# Spectrum of Neandertal introgression across modern-day humans indicates multiple episodes of human-Neandertal interbreeding

**DOI:** 10.1101/343087

**Authors:** Fernando A. Villanea, Joshua G. Schraiber

## Abstract

Neandertals and anatomically modern humans overlapped geographically for a period of over 30,000 years following human migration out of Africa. During this period, Neandertals and humans interbred, as evidenced by Neandertal portions of the genome carried by non-African individuals today. A key observation is that the proportion of Neandertal ancestry is ∼12-20% higher in East Asian individuals relative to European individuals. Here, we explore various demographic models that could explain this observation. These include distinguishing between a single admixture event and multiple Neandertal contributions to either population, and the hypothesis that reduced Neandertal ancestry in modern Europeans resulted from more recent admixture with a ghost population that lacked a Neandertal ancestry component (the “dilution” hypothesis). In order to summarize the asymmetric pattern of Neandertal allele frequencies, we compile the joint fragment frequency spectrum (FFS) of European and East Asian Neandertal fragments and compare it to both analytical theory and data simulated under various models of admixture. Using maximum likelihood and machine learning, we found that a simple model of a single admixture does not fit the empirical data, and instead favor a model of multiple episodes of gene flow into both European and East Asian populations. These findings indicate more long-term, complex interaction between humans and Neandertals than previously appreciated.

## 2 Introduction

When anatomically modern humans dispersed out of Africa they encountered and hybridized with Neandertals [Green et al., 2010]. The Neandertal component of the modern human genome is ubiquitous in non-African populations, and yet is quantitatively small, representing on average only ∼2% of those genomes [Green et al., 2010, Prüfer et al., 2017]. This pattern of Neandertal ancestry in modern human genomes was initially interpreted as evidence of a single period of admixture, occurring shortly after the out-of-Africa bottleneck [Green et al., 2010, Sankararaman et al., 2012]. However, subsequent research showed that Neandertal ancestry is higher by ∼12-20% in modern East Asian individuals relative to modern European individuals [Prüfer et al., 2017, Meyer et al., 2012, Wall et al., 2013].

Neandertals occupied a vast area of Asia and Europe at the time AMH dispersed outside of Africa (∼75,000 BP [Karmin et al., 2015]), and later Europe and Asia (∼47-55,000 BP [Poznik et al., 2016, Skoglund and Mathieson, 2018]). Moreover, the breakdown of Neandertal segments in modern human genomes is indicative of a time-frame for admixture of 50,000-60,000 BP [Sankararaman et al., 2012, Skoglund and Mathieson, 2018], prior to the diversification of East Asian and European lineages. The genome of Ust’-Ishim, an ancient individual of equidistant relation to modern East Asians and Europeans, has similar levels of Neandertal ancestry as modern Eurasians, but found in longer segments, consistent with an admixture episode occurring ∼52,000-58,000 BP [Fu et al., 2015]. Given the extensive support for a single, shared admixture among Eurasians, there is extensive debate surrounding the observation of increased Neandertal ancestry in East Asians.

There are several hypotheses that may explain the discrepancy in Neandertal ancestry between Europeans and East Asians. It is possible that admixture occurred in a single episode, or ‘pulse’, of gene flow, but demographic and/or selective forces shifted the remaining Neandertal alleles into the frequencies we see in modern populations. Among these explanations are differential strength of purifying selection across Eurasia [Sankararaman et al., 2014] and that modern Europeans lost part of their Neandertal ancestry through ‘dilution’ through a later contribution from a ghost population which had remained unadmixed [Vernot and Akey, 2015, Lazaridis et al., 2016]. It is also possible that admixture occurred multiple times; the first pulse of Neandertal gene flow into the population ancestral to East Asians and Europeans was supplemented by additional pulses after both populations had diverged [Vernot and Akey, 2015, Vernot et al., 2016].

Sankararaman et al. [2014] proposed that differences in the level of Neandertal ancestry in East Asian individuals could be explained by their lower ancestral effective population size relative to Europeans, which would reduce the efficacy of purifying selection against deleterious Neandertal alleles [Harris and Nielsen, 2016]. However, Kim and Lohmueller [2015] found that differences in the strength of purifying selection and population size are unlikely to explain the enrichment of Neandertal ancestry in East Asian individuals. This conclusion was further strengthened by Juric et al. [2016].

Another hypothesis consistent with a single episode of gene flow is that Neandertal ancestry in modern Europeans was diluted by one of the populations that mixed to create modern Europeans [Lazaridis et al., 2014, 2016]. This population, dubbed ‘Basal Eurasian’, possibly migrated out of Africa separately from the population receiving the pulse of Neandertal gene flow, and thus had little to no Neandertal ancestry.

On the other hand, admixture may have occurred multiple times; the first pulse of Neandertal gene flow into the population ancestral to East Asians and Europeans was supplemented by additional episodes after both populations had diverged [Vernot and Akey, 2015, Vernot et al., 2016]. The finding of an individual from Peştera cu Oase, Romania with a recent Neandertal ancestor provides direct evidence of additional episodes of interbreeding, although this individual is unlikely to have contributed to modern-day diversity [Fu et al., 2015]. However, Neandertal ancestry has remained relatively constant across tens of thousands of years of Eurasian history [Petr et al., 2018], suggesting that any additional admixture events must have been smaller scale than the initial episodes of interbreeding.

Here, we study the the asymmetry in the pattern of Neandertal introgression in modern human genomes between individuals of East Asian and European ancestry. We summarize the asymmetric distribution of Neandertal ancestry tracts in the East Asian and European individuals in the 1000 Genomes Project panel in a joint fragment frequency spectrum (FFS) matrix. We first fit analytical models using maximum likelihood to explain the distribution of fragments in European and Asian individuals marginally. Then, we compare the joint FFS to the output of genomic data simulated under specific models of admixture between Neandertals and AMH to achieve a higher resolution picture of the interplay of different demographic forces. Our results support a complex model of admixture, with early admixture occurring before the diversification of European and East Asian lineages, and secondary episodes of gene flow into both populations independently.

## 3 Results

We constructed the joint fragment frequency spectrum by analyzing published datasets of Neandertal fragment calls in 1000 genomes individuals [Sankararaman et al., 2014, Steinrücken et al., 2018]. To avoid complications due to partially overlapping fragments and difficulties calling the edges of fragments, we computed fragment frequencies by sampling a single site every 100kb and asking how many haplotypes were introgressed with confidence above a certain cutoff (Figure 1). Our main results make use fragments called with posterior probability of 0.45 in the Steinrücken dataset, although we verified robustness across a range of cutoffs and between datasets (Supplementary Material). The observed average proportion of Neandertal ancestry in European individuals was 0.0137 and 0.0164 in East Asian individuals, corresponding to an average enrichment of 19.6% in East Asian individuals (Supplementary Figure 5 shows how this quantity changes across cutoffs).

**Figure 1:**
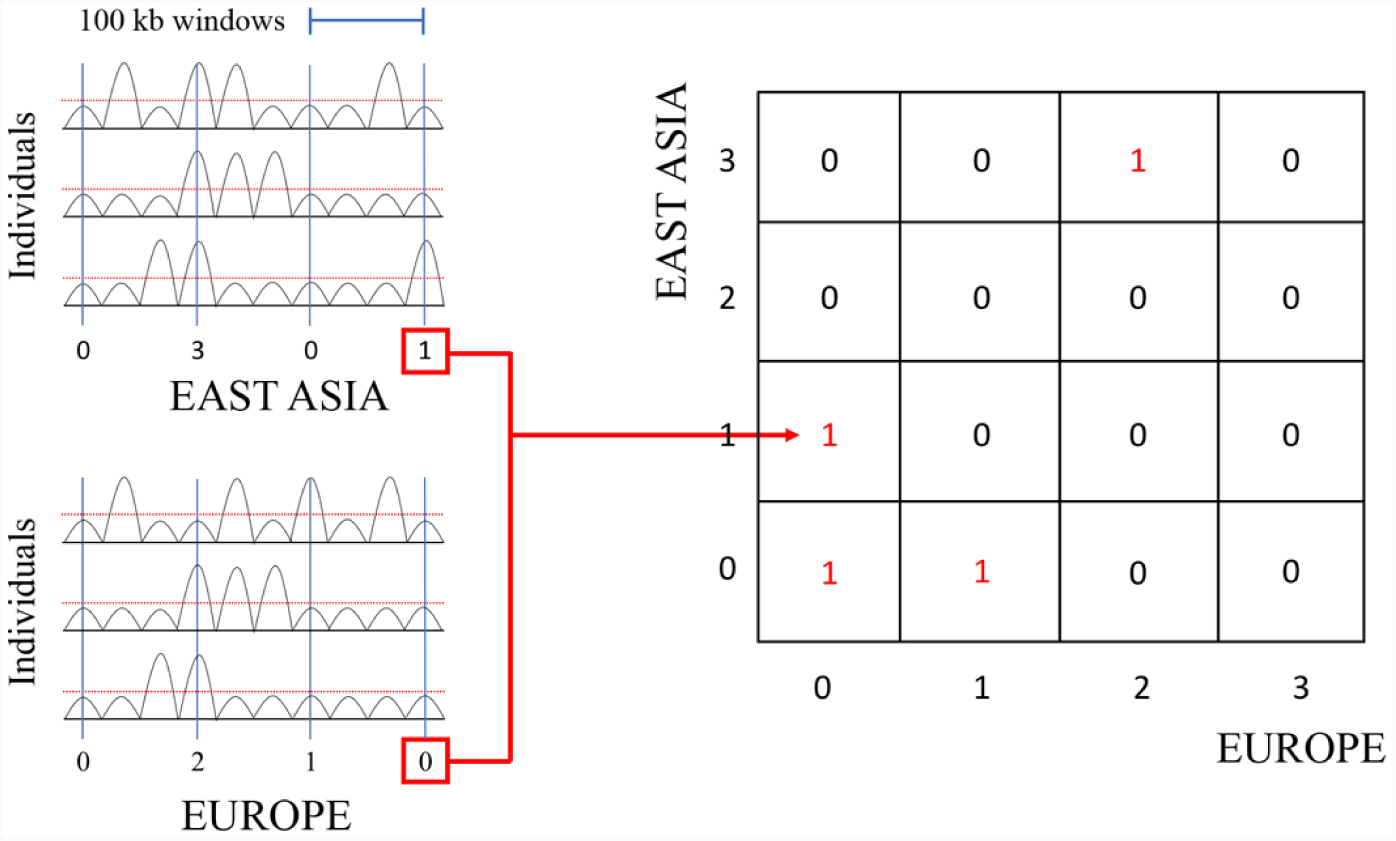
Graphic representation of the process to make an FFS based on the posterior probability calls from Steinrücken et al. [2018]. At a single position every 100 kb, introgression is assigned for individuals presenting a posterior probability above the global cut-off. The count of individuals between East Asian and European samples determines the cell in the FFS where that SNP is counted

We first developed analytic theory to understand what the FFS would look like in each population separately under different demographic models. To our surprise, we found that when looking only at the marginal distribution of introgressed fragment frequencies, the one pulse model and the dilution model are not statistically identifiable (Supplementary Material). On the other hand, the two pulse model is identifiable. Moreover, the analytic theory reveals that population size history only impacts the FFS within each population as a function of effective population size; intuitively, this arises because once fragments enter the population, their frequency dynamics only depend on the effective population size, rather than the specifics of the population size history. With this in mind, we developed a maximum likelihood procedure to fit a one pulse and a two pulse model to the European and East Asian marginal spectra (Methods), and found strong support for the two pulse model in both cases (Λ = 193.91 in East Asians, nominal *p* = 7 × 10^−43^; Λ = 212.64 in Europeans, nomial *p* = 6 × 10^−47^. Figure 2a and b). A subsequent goodness-of-fit test strongly rejected the fit of the one pulse model (*p* = 2 × 10^−26^ in East Asians, *p* = 0.0 in Europeans; *χ*^2^ goodness of fit test) but could not reject the fit of the two pulse model in either populations (*p* = 1 in East Asians, *p* = 0.95 in Europe; *χ*^2^ goodness of fit test); see also Supplementary Figure 1, which shows the residuals of each fit. Thus, we concluded from analyzing each population in isolation that that the history of admixture was complex, and involved multiple matings with Neandertals.

**Figure 2:**
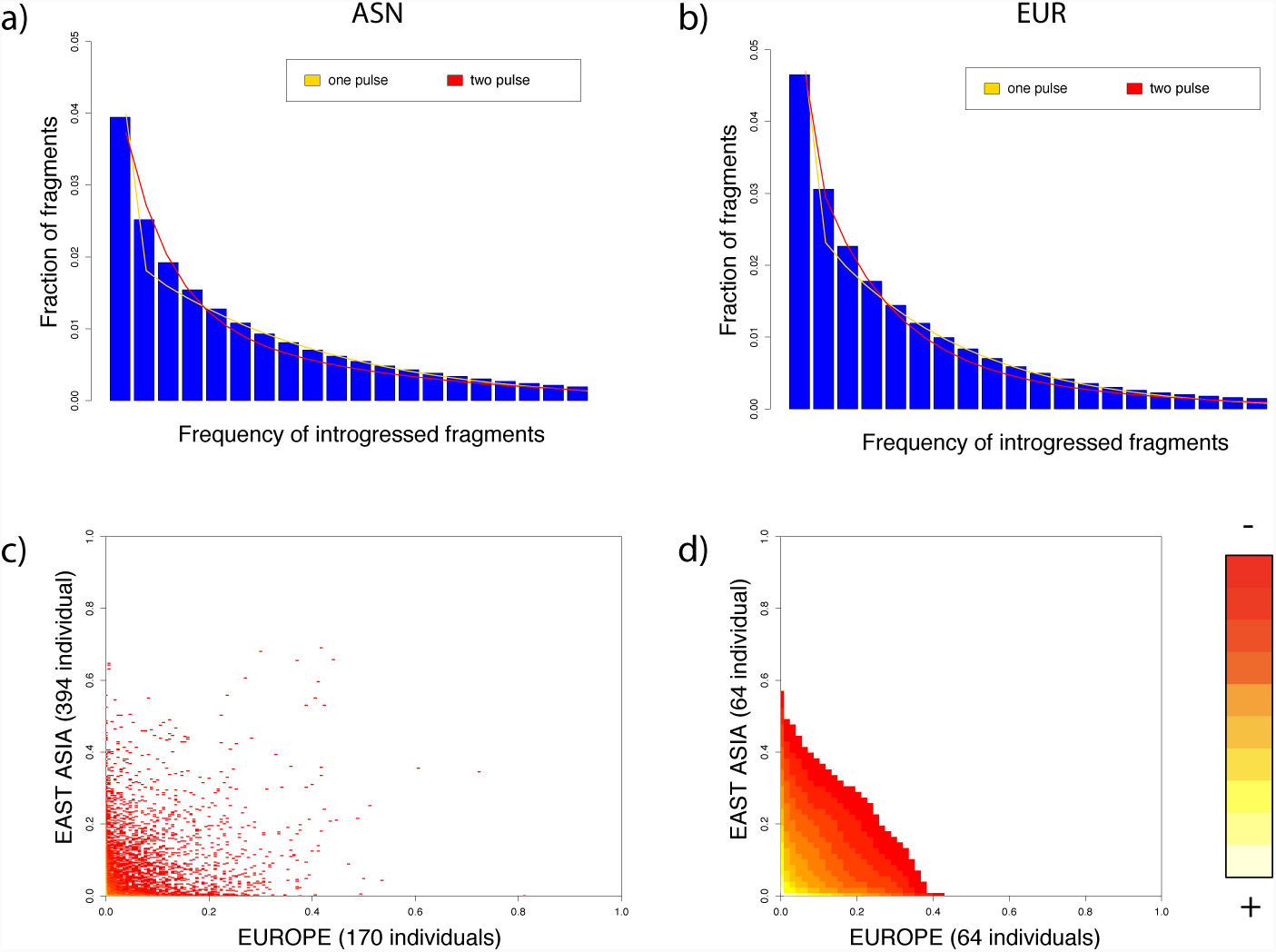
a,b) Marginal SFS for the East Asian and European populations (first 20 bins, excluding the 0 bin). The lines represent the best fitted one pulse and two pulse model for each population. c) FFS of the Steinrücken et al. [2018] introgression data. d)FFS of the Steinrücken et al. [2018] introgression data projected down to 64×64 bins, as used to train the FCNN.

Nonetheless, looking at each population individually, we did not have power to estimate the relative contribution of dilution and multiple admixtures in shaping the patterns of Neandertal fragments seen between Europe and Asia. To gain a more global picture of the history of human-Neandertal interbreeding, we developed a supervised machine learning approach. A difficulty simulating Neandertal admixture is the large number of free parameters associated with modeling multiple populations from which we have incomplete demographic information. Supervised machine learning applied to genomic datasets is becoming a popular solution for inference [for examples see: Schrider and Kern, 2016, Ronen et al., 2013, Sheehan and Song, 2016]. Of particular interest to this study, supervised machine learning has demonstrated the capacity for optimizing the predictive accuracy of an algorithm in datasets that cannot be adequately modeled with a reasonable number of parameters [Schrider and Kern, 2018]. In practice, this results in the ability to describe natural processes even based on incomplete or imprecise models [Schrider and Kern, 2018]. Supervised machine learning implementing hidden layers, or deep learning, is particularly effective in population genetic inference and learning informative features of data [Sheehan and Song, 2016]. A definitive advantage of deep learning is how it makes full use of datasets to learn the mapping of data to parameters, allowing inference from sparse data sets [Schrider and Kern, 2018]. Comparable likelihood-free inference methods, such as ABC, typically use a rejection algorithm, resulting in most simulations being thrown away. This necessitates a very large number of simulations for accurate inference [Sheehan and Song, 2016, Bengio et al., 2007]. Deep learning methods also have the potential to generalize in non-local ways, allowing them to make predictions for data not covered by the training set [Bengio et al., 2007, Schrider and Kern, 2018].

We simulated Neandertal admixture by specifying five demographic models with different numbers of admixture events (Figure 3), and produce FFS under a wide range of parameters. We used the simulated FFS to train a fully-connected neural network (FCNN). The trained network classified models successfully ∼58% of the time, well above the 20% expected by chance, and was not overfit to the training data (Supplementary Figure 2). We then examined how the precision of the prediction changed when we required different levels of support for the chosen model (Figure 4a). Crucially, we see that when the classifier has high confidence in a prediction, it is very often correct, and that multiple pulse models are not often confused the dilution model (Supplementary Figure 3).

**Figure 3:**
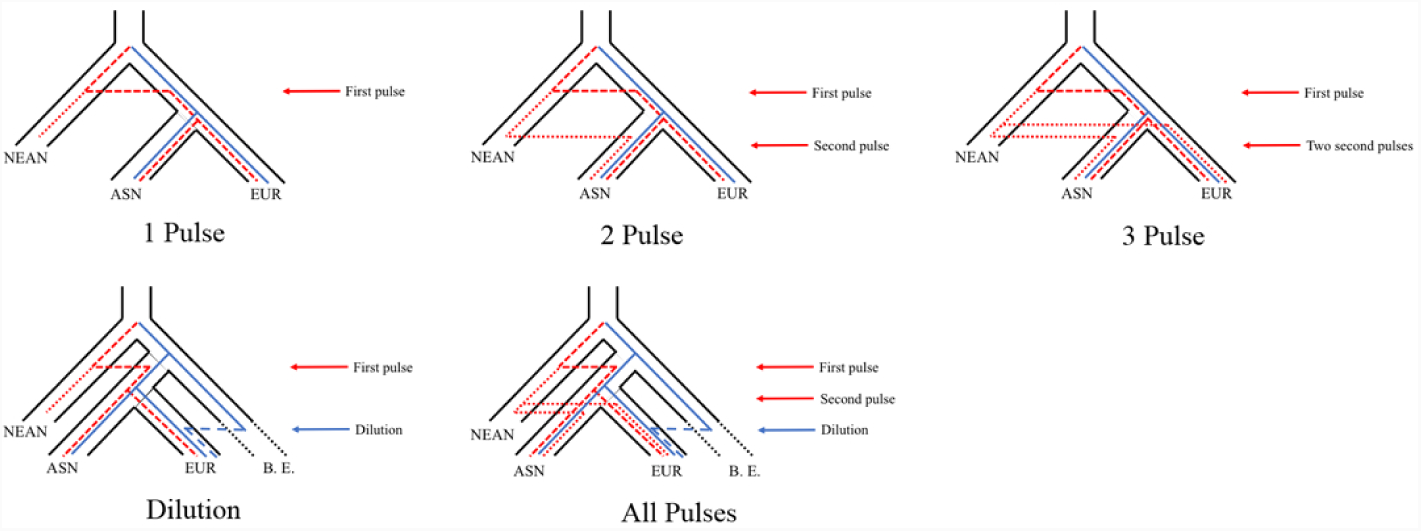
Representation of the five different demographic models simulated in `msprime.` Lines within each population phylogeny indicate different paths that alleles could enter into modern populations, with red lines indicating Neandertal alleles and blue lines indicating non-Neandertal alleles.

**Figure 4:**
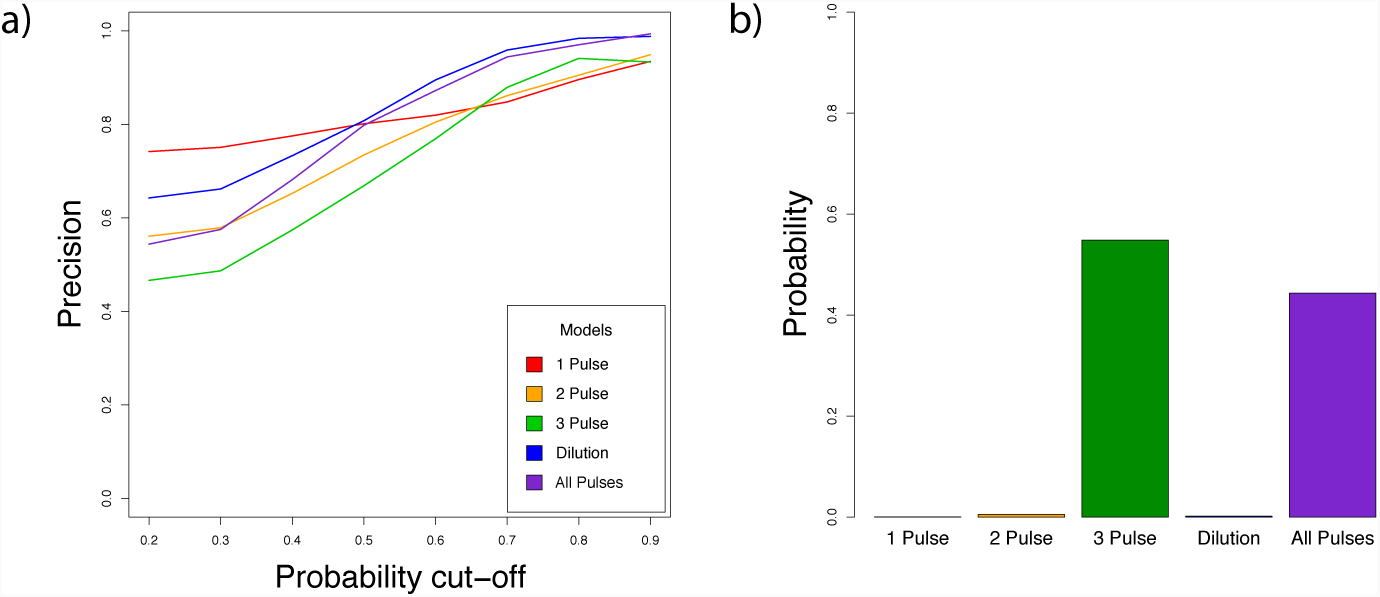
(a) Posterior probability that a chosen model is correct (precision), for all models under different levels of support for the chosen model. The x axis shows the probability cutoff that we used to classify models, and the y axis shows the precision. Each line corresponds to a different model. Simulated datasets in which no model surpassed the cutoff were deemed unclassified. (b) Posterior probability of the empirical introgression data matching each of the five demographic models, determined by the FCNN classifier.

Finally, we applied the trained FCNN to our empirical joint FFS (Figure 4b). Strikingly, we found that the FCNN supported our two most complicated demographic models, favoring a model with 3 pulses of admixture (Posterior probability ∼ 0.55), and with a lower probability, a model with 3 pulses of admixture and dilution (Posterior probability ∼ 0.44). These results are consistent across a range of cutoffs for calling introgressed fragments (Supplementary Figure 4), are robust to errors in fragment calling (Supplementary Figures 9 and 10), and dovetail with our maximum likelihood results showing that the best fit model must include multiple episodes of human-Neandertal interbreeding.

## 4 Discussion

Despite initial indications of a simple history of admixture between humans and Neandertals, more detailed analyses suggested that there might be additional, population specific episodes of admixture. By analyzing the joint fragment frequency spectrum of introgressed Neandertal haplotypes in modern Europeans and Asians, we found strong support for a model of multiple admixture events. Specifically, our results support a model in which the original pulse of introgression into the ancestral Eurasian population is supplemented with additional pulses to both European and East Asian populations after those populations diverge, resulting in elevated Neandertal ancestry in East Asians relative to Europeans. This is similar to a model recently proposed by Vernot et al. [2016] for explanation differential levels of Neandertal ancestry across Europe, Asia, and Melanesia. Importantly, our results exclude a demographic model where the difference in Neandertal ancestry between Europeans and East Asians is driven primarily through dilution of Neandertal ancestry in Europe due to recent admixture with Basal Eurasians, a population lacking Neandertal ancestry. Nonetheless, we cannot exclude dilution as playing a role in the differences in Neandertal ancestry between Europe and East Asia; a model which includes multiple pulses of Neandertal introgression and dilution through Basal Eurasians was the second likeliest model in the five model comparison. Given the evidence that Basal Eurasians contributed to the modern European gene pool [Lazaridis et al., 2016], we suspect that dilution does play a role in shaping the pattern of Neandertal ancestry across Eurasia. However, a large amount of dilution would be necessary if it were the only factor explaining the ∼19.6% difference in Neandertal ancestry between Europe and East Asia, in contrast with recent work that inferred a smaller (∼9.4%) contribution of Basal Eurasians to modern European individuals [Kamm et al., 2018].

Several confounding factors could impact our inference. Although it is unlikely that differential purifying selection is responsible for the discrepancy between European and East Asian Neandertal ancestry [Kim and Lohmueller, 2015, Vernot and Akey, 2015, Juric et al., 2016], some Neandertal ancestry was likely deleterious [Harris and Nielsen, 2016, Juric et al., 2016] and our models assume neutrality. However, the strength of selection against introgressed fragments is likely to be small compared to the demographic forces at work; moreover, there is relatively little evidence of strong differences in the strength of selection between different non-African populations [Simons et al., 2014, Do et al., 2015]. To explore the impact of selection, we obtained the FFS from simulations of deleterious Neandertal ancestry by Petr et al. [2018] and asked if we classified their scenarios with selection as a two pulse model using maximum likelihood. We found that we rejected a one pulse model at the 5% level in only 1 out of 15 different simulations with selection, suggesting that we are not likely to misclassify selection against Neandertal ancestry as a two pulse model.

Of additional concern is power to detect fragments in each population. To address this, we implemented a model of fragment calling errors (Supplementary Material). Based on simulations done by Steinrücken et al. [2018], we expect false positive rates of approximately .1% and false negative rates of approximate 1%; such rates do not cause substantial shifts in the FFS (Supplementary Figure 6). Moreover, after extensive simulations, we found that the neural network trained with errors is robust to false positive fragment calls at a rate of 0.2%, and produce consistent results when applied to the real data (Supplementary Material). Finally, in an attempt to see inside the “black-box” of the fully connected network, we examined how the weights propagate from each entry of the JFFS to the final assignments (Supplementary Material). In doing so, we found that moderate frequency haplotypes are most important to distinguish between models (Supplementary Figure 7), whereas errors in calling fragments are most likely to impact low frequency haplotypes. This, combined with the fact that our results are robust across two different datasets and a range of cutoffs for determining archaic ancestry, convince us that our results are robust to errors in fragment calling.

In addition, it is possible that some of the Neandertal ancestry in East Asia has been misclassified, and in fact originated from Denisovan introgression, mimicking the signal of additional pulses of Neandertal introgression. To address this concern, inferred the position of Denisova fragments based on data from [Browning et al., 2018] to mask 1.6% of positions in the genome (Methods). This resulted in 0.49% of Neandertal introgressed sites being removed from the Steinrücken et al. [2018] data used in all analyses. These problems are likely to be further resolved as our ability to make accurate introgression calls for the various ancient human populations improves in the future.

Our work provides additional evidence for the ubiquity of archaic admixture in recent human history, consistent with recent work shows that humans mixed with Denisovans multiple times [Browning et al., 2018]. Though we find that additional pulses of admixture in both East Asians and Europeans are necessary to explain the distribution of Neandertal ancestry in Eurasia, we are unable to settle *why* East Asians have elevated Neandertal ancestry. Interestingly, in contrast to Denisovans, there does not seem to be evidence of Neandertal population structure within introgressed fragments [Browning et al., 2018]. Combined with our results, this indicates that the Neandertal population or populations that admixed with Eurasians must have been relatively closely related. This is consistent with the established inference of a long-term small effective size across Neandertals [Prüfer et al., 2014, 2017], which has held up to scrutiny despite some claims of a larger Neandertal effective size [Rogers et al., 2017a,b, Mafessoni and Prüfer, 2017]. Thus, we believe that a likely explanation for our results is that gene flow between humans and Neandertals was intermittent and ongoing, but in a somewhat geographically restricted region. Thus, differential levels of admixture between different Eurasian groups may primarily reflect how long those populations coexisted with Neandertals in that region.

## 5 Methods

### 5.1 Data

We obtained the joint fragment frequency spectrum by first downloading publicly available Neandertal introgression calls from two sources [Sankararaman et al., 2014, Steinrücken et al., 2018]. The Sankararaman data consists of the location of introgressed fragments along the genome in each phased haplotype for the 1000 Genomes Project populations. The Steinrücken data consists of the probability of Neandertal origin in 500 bp windows of the genome across each phased haplotype for the Central European (CEU) and Han Chinese (CHB) and Southern Han Chinese (CHS) from the 1000 Genomes Project. We computed fragment frequencies by sampling a single site every 100kb, from both sources of data. To compute the Joint Fragment Frequency Spectrum (FFS), we counted how many haplotypes were introgressed at each position. For the Steinrücken data, we called a site introgressed if it has a posterior probability of being introgressed above 45%. We then applied the 1000 Genomes accessibility mask (downloaded from ftp://ftp.1000genomes.ebi.ac.uk/vol1/ftp/release/20130502/supporting/accessible_genome_masks/20141020.pilot_mask.whole_genome.bed). In the supplement, we show that our results are robust to the cutoff and consistent between both datasets. We also masked Denisova fragments that were falsely called as Neandertal by downloading the S’ fragment calls from Browning et al. [2018] and masking any fragment that matched Denisova > 35% and Neandertal < 25%. Finally, we masked the (0,0), (0,1), (1,0), and (1,1) position of the FFS matrix, in order to reduce the impact of false negative and false positive fragment calls.

### 5.2 Analytical Model

We model introgression of intensity *f* as injection of alleles at frequency *f* into the population at the time of introgression. In the one pulse model, this results in an exact expression for the expected fragment frequency spectrum under the Wright-Fisher diffusion model (Supplementary Material). Multiple pulse models could be solved analytically using a dynamic programming algorithm as in Kamm et al. [2018], but we instead approximate the expected frequency spectrum by making the approximation that the probability of sampling *k* introgressed haplotypes in a sample of size *n* + 1 is the same as sampling *k* haplotypes in a sample of size *n* for large *n* (c.f. Jouganous et al. [2017]). This results in closed-form expressions for the expected frequency spectrum under both the two-pulse and the dilution model (Supplementary Material). With an expected frequency spectrum given parameters *θ* and model *ℳ, p*_*n,k*_(*θ, ℳ*) = ℙ (k out of n haplotypes are introgressed|*θ, ℳ*), we compute the likelihood

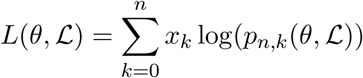

where *x*_*k*_ is the number of fragments found in *k* out of *n* individuals. We optimized the likelihood using `scipy,` and compared models using the likelihood ratio statistic

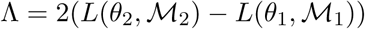

where *θ*_*i*_ and *ℳ*_*i*_ correspond to the *i*-pulse model. Under the null, Λ should be *χ*^2^ distributed with 2 degrees of freedom (since there are 2 additional parameters in the two pulse model). Simulations in the Supplementary Material suggest that p-values under this model are well calibrated, despite the impact of linkage; thus, we opted to report nominal p-values from our likelihood ratio test.

### 5.3 Simulations

We used `msprime` [Kelleher et al., 2016], to simulate Neandertal introgression into two modern populations with multiple potential admixture episodes and dilution from Basal Eurasians (Figure 3). For each replicate, we simulated the complete genomes for 170 European individuals, and 394 East Asian individuals, matching the sampling available from the 1000 Genomes Project panel. We used the human recombination map (downloaded from http://www.well.ox.ac.uk/∼anjali/AAmap/; Hinch et al. [2011]). In each simulation we mimicked our sampling scheme on the real data by sampling 1 site every 100kb and calling a Neandertal fragment by asking which individuals coalesced with the Neandertal sample more recently than the human-Neandertal population split time.

For each simulation, we drew demographic parameters, including effective population sizes and divergence times, from uniform distributions. For effective population sizes we used 500-5000 individuals for Neandertals, 5000-50000 for Eurasians, 5000-100000 individuals for the European and East Asian populations. For divergence times, we used 26000-12000 generations for Neandertals and humans, and 1300-2000 generations for the Eurasian split. The divergence between Basal Eurasians and Eurasians was fixed at 3000 generations. Lastly, we drew introgression times between 1500-3000 generations for gene flow into Eurasians, 800-2000 generations for gene flow into the European and East Asian populations. The time for the introgression event between Basal Eurasians and Europeans (dilution) was drawn from an uniform distribution of 200-2000 generations.

In order to ensure that our simulations focused on the correct parameter space, we constrained the resulting amount of Neandertal introgression in the modern European and East Asian genomes. The average Neandertal ancestry *a* was drawn from an uniform distribution between 0.01-0.03, and the difference in ancestry *d* between the East Asian and European populations was drawn from an uniform distribution between 0-0.01. We then determined the introgression intensity given *a* and *d* (Supplementary Material).

Our simulation pipeline is available at https://github.com/Villanea/Neandertal_admix/blob/master/n_admix_10.py, which generates the simulated genomes, identifies Neandertal introgression in 100kb windows, applies the 1000 Genomes accessibility mask, and outputs a FFS and accompanying parameters for each replicate.

### 5.4 Machine Learning (FCNN)

Using the resulting joint FFS, we trained a simple fully-connected neural network (FCNN) to categorize a joint FFS into one of five demographic models. The network was implemented in Keras [Chollet et al., 2015] using a TensorFlow back-end. The network used a simple architecture of three Dense layers (from 1024 nodes, to 512 nodes, to 64 nodes), each followed by a dropout layer (0.20). The code used to implement the network can be found at https://github.com/Villanea/Neandertal_admix/blob/master/Fully_connected_network.py.

## 6 Acknowledgments

We are grateful to Sara Mathieson and Jeff Spence for several useful discussions about neural network architecture and appropriate methods for training neural networks. We also wish to thank Jeff Spence and Matthias Steinrücken for extensive discussions on errors in fragment calling. Iain Mathieson, Sara Mathieson, and Jeff Spence provided invaluable feedback on an early draft of this manuscript that helped improve its clarity. Kelley Harris provided invaluable discussions during the conception and work of this manuscript. We’re grateful to Martin Petr and Benjamin Vernot for sharing processed simulation data with us, and for disucssions about the impact of selection on Neandertal ancestry. This work was supported by NIH grant R35 GM124745.

